# Loss of m6A unmasks a senescence-associated inflammatory program in human microglia

**DOI:** 10.64898/2026.07.06.736874

**Authors:** Deepak Pathak, Rashmi Basavaraj, Lina Sun, Yi-Lan Weng

## Abstract

Chronic microglial inflammation and cellular senescence are hallmarks of the aging brain, yet the molecular events that lock microglia into durable inflammatory states remain poorly understood. Prior studies using genetic manipulation of METTL3 have implicated the RNA modification N6-methyladenosine (m6A) in senescence. However, because METTL3 also has m6A-independent functions, loss-of-function approaches cannot distinguish whether senescence arises from reduced m6A itself or from broader disruption of METTL3-dependent pathways. This question is further complicated in microglia, where METTL3 has been reported to promote acute inflammatory activation, suggesting that m6A may have context-dependent effects on immune state. Whether sustained reduction of m6A is sufficient to drive microglial senescence has therefore remained unresolved.

Here, we show that selective catalytic inhibition of METTL3 with STM2457 lowers global m6A and is sufficient to induce a senescence-like inflammatory state in human HMC3 microglia. This state includes increased senescence-associated β-galactosidase activity, elongated cellular morphology, reduced proliferation, Lamin B1 loss, and remodeling of nuclear architecture. Transcriptomic profiling revealed suppression of mitotic gene programs together with a SASP-like inflammatory output marked by NF-κB and interferon signatures. Rather than causing broad transposable-element derepression, m6A inhibition promoted cytoplasmic double-stranded RNA accumulation and a selective HERVK-associated response. These findings support a model in which m6A helps preserve microglial homeostasis by limiting immunogenic RNA accumulation and senescence-associated inflammatory remodeling. Together, our study identifies m6A as a safeguard against microglial senescence and suggests that reduced m6A-dependent RNA regulation may contribute to chronic inflammatory remodeling in the aging brain.

## Introduction

Chronic, low-grade inflammation is a defining feature of the aging brain and a major contributor to neurodegenerative disease. Microglia, the resident innate immune cells of the central nervous system, continuously survey the brain parenchyma and respond to tissue damage, neuronal dysfunction, and protein aggregation by clearing cellular debris and supporting neuronal homeostasis. With age, however, a subset of microglia lose this homeostatic, surveillant state and acquire persistent inflammatory phenotypes. Aged microglia increasingly display features of cellular senescence, including reduced proliferative capacity, altered nuclear organization, and sustained production of senescence-associated secretory phenotype (SASP) factors^1–5^. Although these states are increasingly recognized as contributors to brain aging and disease, the upstream molecular events that convert homeostatic microglia into durably inflammatory, senescence-like cells remain poorly defined.

N6-methyladenosine (m6A) is the most abundant internal modification of eukaryotic mRNA and plays broad roles in post-transcriptional gene regulation^6^. Installed primarily by the METTL3/METTL14 methyltransferase complex and decoded by m6A reader proteins, m6A influences RNA splicing, export, translation, and decay^6–8^. Emerging evidence indicates that m6A function extends beyond protein-coding transcripts to regulate repeat-derived RNAs, particularly retrotransposon-derived transcripts such as LINE-1 and endogenous retroviruses (ERVs)^9–11^. Loss of this regulatory layer may permit repeat-derived or structured RNAs to accumulate, alter their subcellular handling, and generate immunostimulatory nucleic acid species, including dsRNA, that activate innate immune signaling^12–15^. Because ERV reactivation, LINE-1 derepression, and accumulation of immunostimulatory repeat-derived nucleic acids have been implicated in the induction and reinforcement of cellular senescence, m6A-dependent control of these RNA species may be especially important for limiting age-associated inflammatory remodeling^16,17^.

Consistent with this possibility, reduced m6A abundance or impaired METTL3/METTL14 function has been associated with senescence in several cell types^18–20^. However, most of this evidence comes from genetic depletion of writer-complex subunits, which removes both m6A deposition and potential methylation-independent or scaffolding functions of these proteins. Therefore, whether catalytic reduction of m6A itself is sufficient to initiate a coordinated senescence-like inflammatory program remains unresolved.

This question is particularly important in microglia. Prior studies of acute immune challenge or injury models have suggested that m6A signaling can shape microglial inflammatory activation, with METTL3 promoting LPS-induced inflammatory responses and neuroinflammatory gene programs in several settings^21,22^. However, acute inflammatory activation and cellular senescence are distinct biological states. Whether sustained reduction of m6A suppresses inflammatory activation or instead drives a chronic senescence-like program in microglia has not been directly tested.

Here, we test whether catalytic inhibition of METTL3 is sufficient to drive senescence-like remodeling in human HMC3 microglia. Using the selective METTL3 inhibitor STM2457^23^ to reduce m6A without depleting the METTL3/METTL14 complex, we find that m6A reduction induces a coordinated senescence-like state characterized by reduced proliferation, SASP-like NF-κB and interferon inflammatory programs, cytoplasmic dsRNA accumulation, a selective HERV-associated response, and remodeling of nuclear architecture. Together, our findings identify m6A as a safeguard against microglial senescence and suggest that loss of m6A-dependent RNA control may provide one route by which microglia enter chronic inflammatory states associated with brain aging.

## Results

### Catalytic inhibition of METTL3 reduces global m6A and induces a senescence-like state in human HMC3 microglia

To determine whether reduced m6A is sufficient to induce senescence-like phenotypes in human microglia, we treated HMC3 cells with the selective METTL3 catalytic inhibitor STM2457 over a 96-h time course (Figure 1a). m6A dot-blot analysis confirmed a substantial reduction in global m6A levels compared with vehicle-treated cells, indicating effective inhibition of METTL3 methyltransferase activity under these conditions (Figure 1b).

**Figure 1.**
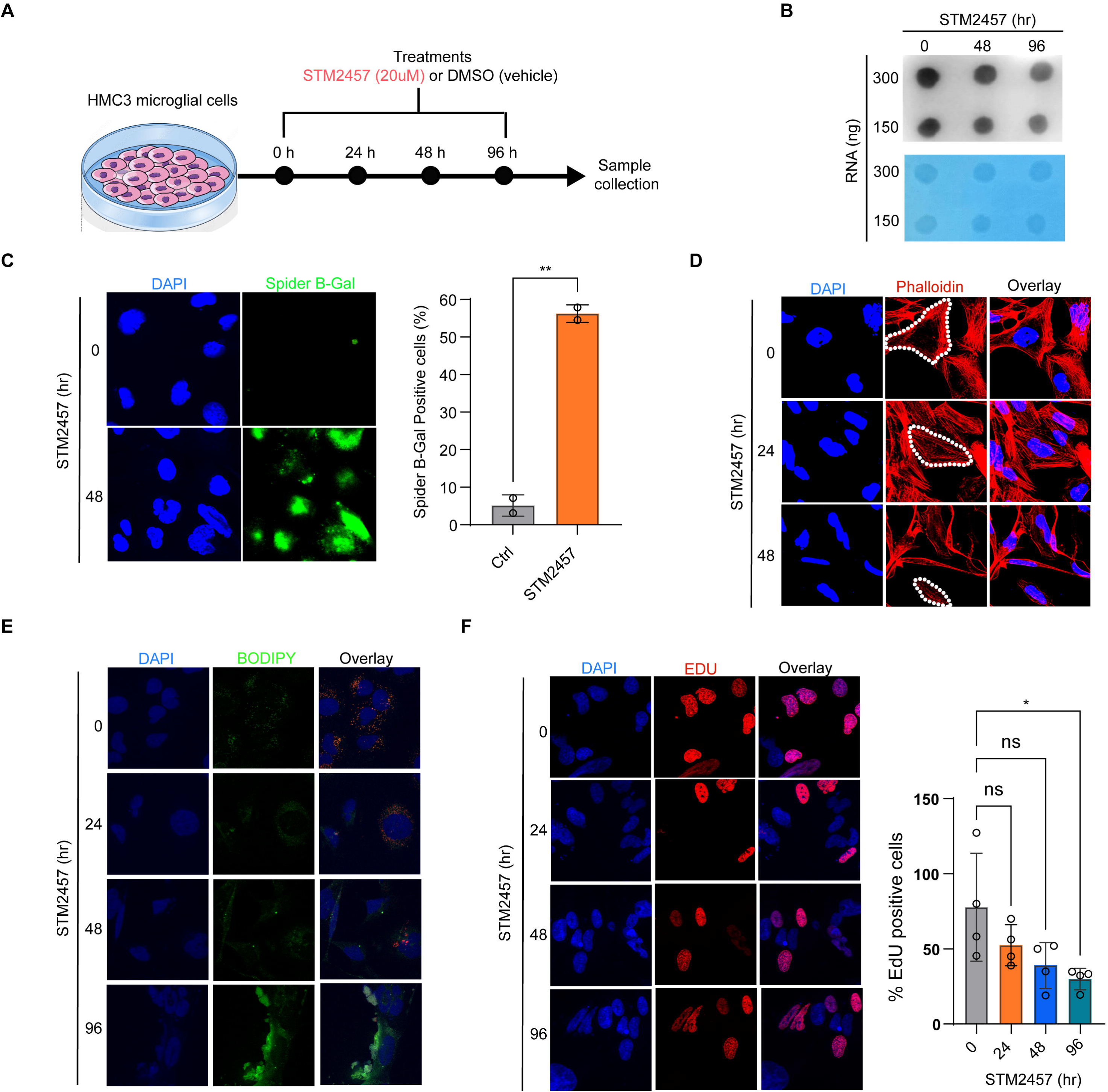
Catalytic inhibition of METTL3 reduces global m6A and induces a senescence-like state in human HMC3 microglia. (a) Schematic of the STM2457 treatment time course (up to 96 h). (b) Representative m6A dot blot showing reduced global m6A in STM2457-treated cells relative to vehicle-treated controls; methylene blue staining indicates total RNA loading. (c) Quantification of senescence-associated β-galactosidase (SA-β-gal)-positive cells following STM2457 or vehicle treatment. (d) Representative images of cell morphology and quantification of mean cell area. (e) BODIPY staining and quantification of intracellular neutral lipid droplets. (f) Quantification of EdU incorporation upon STM2457 96 hr treatment. Data are presented as mean ± SEM from three independent biological experiments, with 5 fields quantified per condition per experiment. Statistical significance was determined by two-tailed unpaired Student’s t test; P values are indicated in the figure.

We next assessed cellular features commonly associated with senescence. Because increased lysosomal β-galactosidase activity is a canonical senescence marker, we first measured senescence-associated β-galactosidase (SA-β-gal) staining. STM2457 markedly increased the proportion of SA-β-gal-positive cells compared with vehicle-treated controls (Figure 1c). Senescence is also accompanied by changes in cell morphology, so we examined cell shape and size. STM2457-treated cells became elongated and thin, with quantification showing a reduced mean cell area (Figure 1d).

Aging and dysfunctional microglial states are often associated with altered lipid metabolism, including the accumulation of intracellular lipid droplets^24,25^. We therefore asked whether m6A inhibition also promotes lipid droplet accumulation. BODIPY 493/503 staining revealed increased neutral lipid droplets in STM2457-treated cells compared with vehicle-treated controls (Figure 1e). Because senescence requires withdrawal from the cell cycle, we then measured proliferation by EdU incorporation. STM2457 progressively reduced the proportion of EdU-positive cells over the treatment time course, indicating impaired proliferation (Figure 1f).

To determine whether these phenotypes reflected senescence-like remodeling rather than nonspecific toxicity, we assessed cell death under the same treatment conditions. STM2457 did not cause substantial cell death (data not shown). Together, these results show that METTL3 catalytic inhibition reduces m6A and induces a senescence-like phenotype in human microglia, characterized by increased SA-β-gal activity, altered morphology, lipid droplet accumulation, and reduced proliferation without overt cytotoxicity.

### m6A inhibition triggers a coordinated growth-arrest and inflammatory program

To define the molecular changes associated with the senescence-like phenotype, we performed RNA sequencing on HMC3 cells treated with STM2457 or vehicle for 48 h using four biological replicates per condition. STM2457 treatment caused broad transcriptional remodeling, with 865 genes upregulated and 590 genes downregulated relative to vehicle-treated controls (fold change ≥ 1.5, adjusted *P* < 0.05; Figure 2a; Supplementary Table 1).

**Figure 2.**
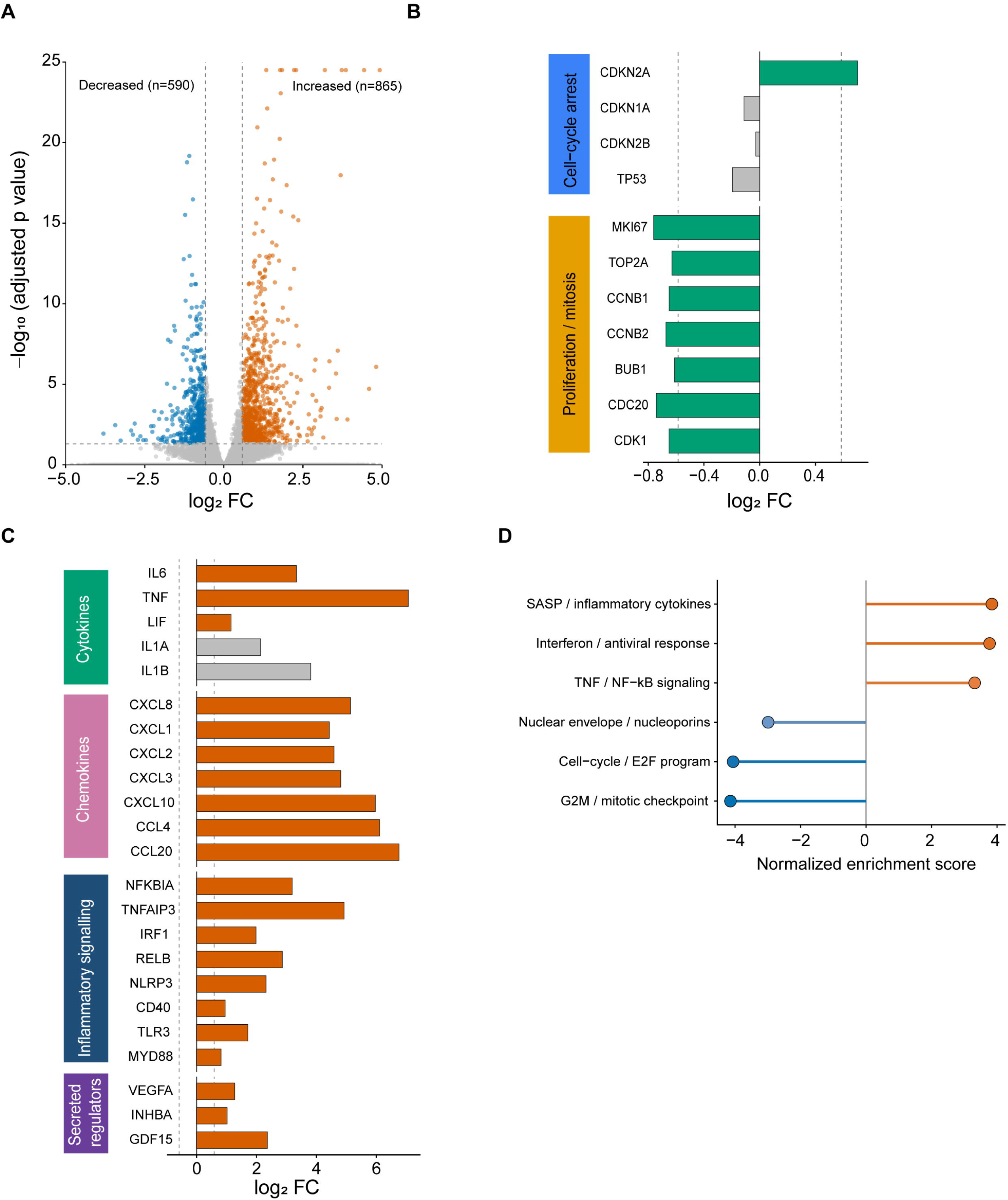
m6A inhibition triggers a coordinated growth-arrest and inflammatory program. (a) Volcano plot of RNA-seq from HMC3 cells treated with STM2457 or vehicle for 48 h (b) Log₂ fold-change (vs vehicle) of representative cell-cycle-arrest and proliferation/mitosis genes; green bars denote significant changes and gray bars non-significant changes. (c) Log₂ fold-change (vs vehicle) of SASP-like inflammatory genes, grouped as cytokines, chemokines, inflammatory-signaling regulators and secreted regulators; orange bars denote significant induction and gray bars non-significant changes. (d) Gene set enrichment analysis (normalized enrichment score) of pathways enriched among STM2457-induced genes (orange) and STM2457-repressed genes (blue).

Consistent with the reduced EdU incorporation observed in Figure 1e, the downregulated gene program was dominated by cell-cycle and mitotic regulators, including *MKI67*, *TOP2A*, *CCNB1*, *CCNB2*, *BUB1*, *CDC20*, and *CDK1* (Figure 2b). In parallel, STM2457 treatment induced *CDKN2A*/p16, a key senescence-associated cell-cycle regulator (Figure 2b). These changes indicate that m6A inhibition suppresses the proliferative and mitotic machinery of HMC3 microglia at the transcriptional level.

Senescent cells often acquire a senescence-associated secretory phenotype (SASP), characterized by sustained expression of inflammatory cytokines, chemokines, and immune regulators ^26,27^. We therefore asked whether m6A inhibition also induced this inflammatory arm of senescence. STM2457 treatment induced a broad SASP-like program that included chemokines such as *CXCL8*, *CXCL1*, *CXCL2*, *CXCL3*, *CXCL10*, *CCL4*, and *CCL20*; cytokines such as *TNF*, *IL6*, and *LIF*; inflammatory regulators including *NLRP3*, *CD40*, *NFKBIA*, *TNFAIP3*, *IRF1*, *RELB*, *TLR3*, and *MYD88*; and secreted regulators including *VEGFA*, *INHBA*, and *GDF15* (Figure 2c). Gene set enrichment analysis confirmed this coordinated transcriptional shift. Genes induced by STM2457 were enriched for SASP and inflammatory cytokine signatures, interferon and antiviral responses, and TNF/NF-κB signaling, whereas downregulated genes were enriched for cell-cycle, G2/M checkpoint, and nuclear envelope and nucleoporin programs (Figure 2d).

Together, these results show that m6A inhibition does not simply reduce proliferation. Instead, it coordinates mitotic shutdown with activation of a SASP-like NF-κB and interferon inflammatory program, defining a transcriptomic state consistent with inflammatory senescence.

### m6A inhibition induces cytoplasmic dsRNA and a selective HERVK-associated response without broad transposable-element derepression

The enrichment of interferon and NF-κB signatures suggested that m6A inhibition may activate innate immune sensing of endogenous nucleic acids. Cytoplasmic double-stranded RNA (dsRNA) is a common trigger of antiviral and inflammatory signaling^28,29^. We therefore asked whether m6A loss promotes dsRNA accumulation. Immunostaining with the dsRNA-specific J2 antibody revealed a marked increase in cytoplasmic dsRNA in STM2457-treated cells compared with vehicle-treated controls (Figure 3a). Thus, METTL3 catalytic inhibition promotes the accumulation of cytoplasmic dsRNA.

**Figure 3.**
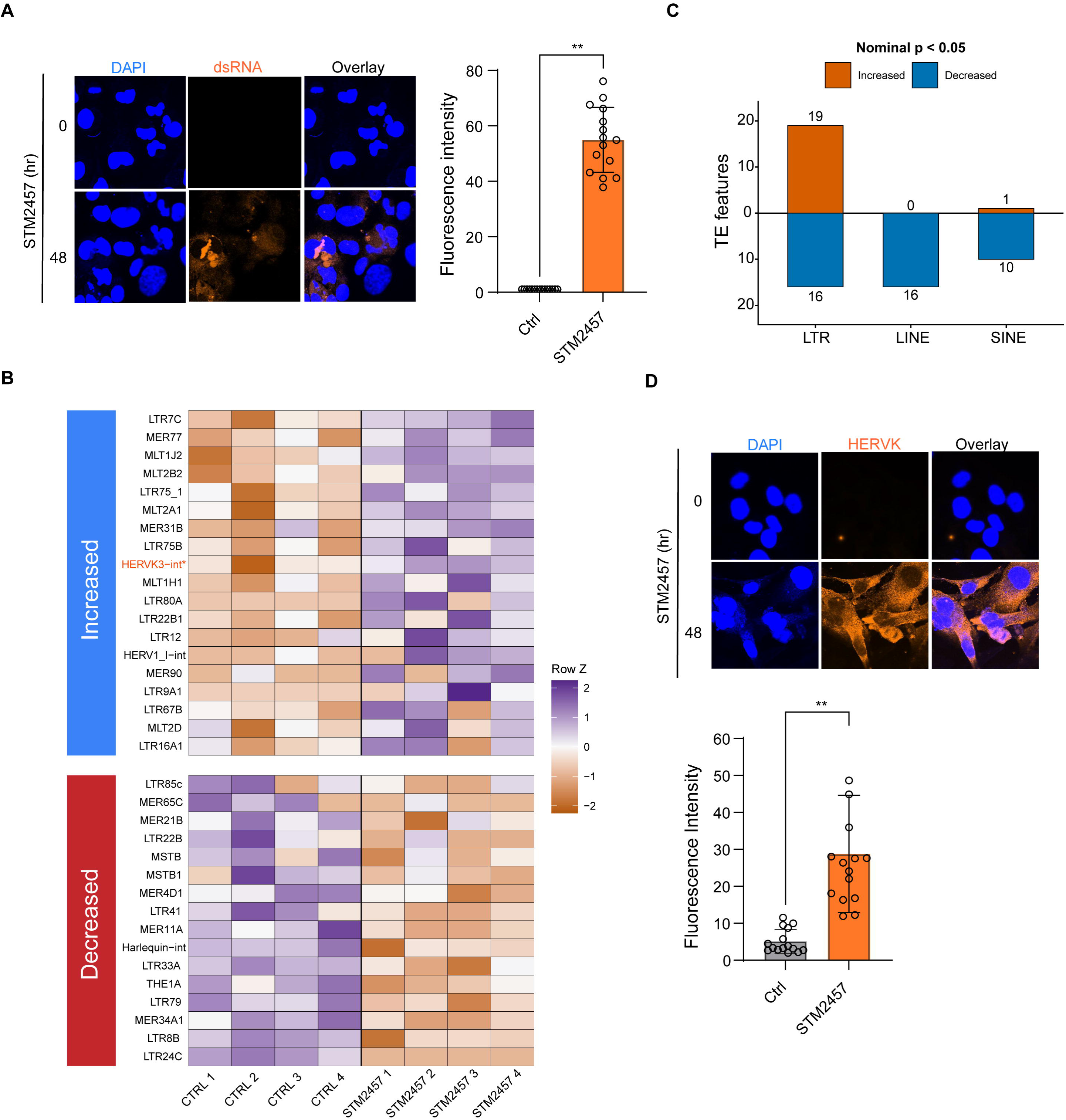
m6A inhibition induces cytoplasmic dsRNA and a selective HERVK-associated response without broad transposable-element derepression. (a) Representative immunofluorescence and quantification of cytoplasmic dsRNA in STM2457- and vehicle-treated HMC3 cells. (b) Differential expression of repeat families in STM2457- versus vehicle-treated cells, quantified with Telescope. (c) TE analysis of LINE, SINE, and LTR repeat classes showing no broad transposable-element activation after STM2457 treatment. (d) Representative immunofluorescence and quantification of HERV envelope (Env) protein in STM2457- and vehicle-treated cells. Data are presented as mean ± SEM; n = 3 independent biological experiments. Statistical significance for the immunofluorescence quantifications was determined by two-tailed unpaired Student’s t test.

We next asked whether repeat-derived RNAs contribute to this dsRNA signal. Repetitive elements are major sources of endogenous immunostimulatory RNA, and their derepression has been linked to sterile inflammation and cellular senescence. LINE-1 activation can drive type I interferon signaling and age-associated inflammation in senescent cells, while endogenous retrovirus reactivation has been proposed to reinforce senescence and tissue aging^16,17^. In addition, m6A has been implicated in the restraint of endogenous retroviral transcripts, as genetic depletion of the METTL3/METTL14 complex in mouse embryonic stem cells increases ERVK and IAP transcript abundance^9^. However, whether catalytic reduction of m6A is sufficient to derepress repetitive elements in human microglia, independent of m6A-independent functions of the writer complex, has not been established.

To test this, we quantified transposable-element and endogenous-retrovirus expression from the 48-h RNA-seq data using two repeat-aware approaches, Telescope and TEtranscripts^30,31^. STM2457 treatment did not produce broad transposable-element activation (Figure 3b). Instead, differential repeat expression was limited to a small number of features, with no global induction of LINE, SINE, or LTR families. TEtranscripts produced concordant results, supporting the conclusion that METTL3 catalytic inhibition does not broadly derepress repetitive elements at this time point (Figure 3c).

Despite the absence of widespread repeat activation, Telescope analysis identified a significant increase in HERVK3-int, a HERVK-related internal repeat feature (Figure 3b). Immunostaining further showed elevated HERV envelope protein in STM2457-treated cells, consistent with activation of a HERV-associated response (Figure 3d). Thus, m6A inhibition induces a selective HERVK-associated signal without broad ERV-family activation.

The increase in cytoplasmic dsRNA was therefore not matched by global repeat derepression. This suggests that the J2-positive signal is unlikely to arise solely from genome-wide transposable-element activation. Instead, it may reflect a more selective and heterogeneous pool of endogenous RNAs, including HERVK-related transcripts, structured host RNAs, antisense transcripts, or retained introns whose abundance, localization, or duplex-forming potential changes when m6A is reduced. Together, these results show that METTL3 catalytic inhibition promotes cytoplasmic dsRNA accumulation together with a restricted HERVK-associated response, identifying immunostimulatory RNA accumulation as a potential feature of m6A-deficient microglia.

### m6A inhibition remodels nuclear architecture and disrupts nuclear protein compartmentalization

Senescence is not only a transcriptional or inflammatory state. It is also accompanied by structural remodeling of the nucleus, including changes in nuclear lamina organization, nuclear shape, and chromatin architecture^32,33^.. Our RNA-seq data suggested that this structural dimension of senescence may be engaged by m6A inhibition. Gene set enrichment analysis revealed downregulation of a nuclear envelope and nuclear pore program in STM2457-treated cells (Figure 2d), including reduced expression of lamina- and nucleoporin-associated genes such as *LMNB1, TMPO, NUP155, NUP107, NUP98, and NUP214*. This raised the possibility that reduced m6A does not merely alter inflammatory and cell-cycle gene expression, but also remodels nuclear structures that support genome organization and RNA processing.

We therefore examined nuclear architecture at the protein and morphological levels. Western blot analysis showed that STM2457 reduced Lamin B1, a core component of the nuclear lamina and a well-established marker of cellular senescence (Figure 4a). Immunofluorescence confirmed decreased Lamin B1 signal in STM2457-treated cells and revealed altered nuclear morphology, including reduced nuclear circularity (Figure 4b). Thus, m6A inhibition converts the transcriptomic signature of nuclear-envelope remodeling into a measurable structural phenotype. Because nuclear lamina and nuclear pore organization are closely linked to nucleocytoplasmic transport, we next asked whether m6A inhibition also affects nuclear protein compartmentalization. We first examined KPNA2, a karyopherin/importin-α family protein involved in nuclear import. In vehicle-treated cells, KPNA2 showed a prominent perinuclear enrichment pattern. After 48 h of STM2457 treatment, this perinuclear pattern was reduced and KPNA2 staining became more diffuse in the cytoplasm. By 96 h, KPNA2 signal was further redistributed outside the nucleus (Figure 4c). These data suggest that m6A inhibition alters the localization pattern of a nuclear import factor.

**Figure 4.**
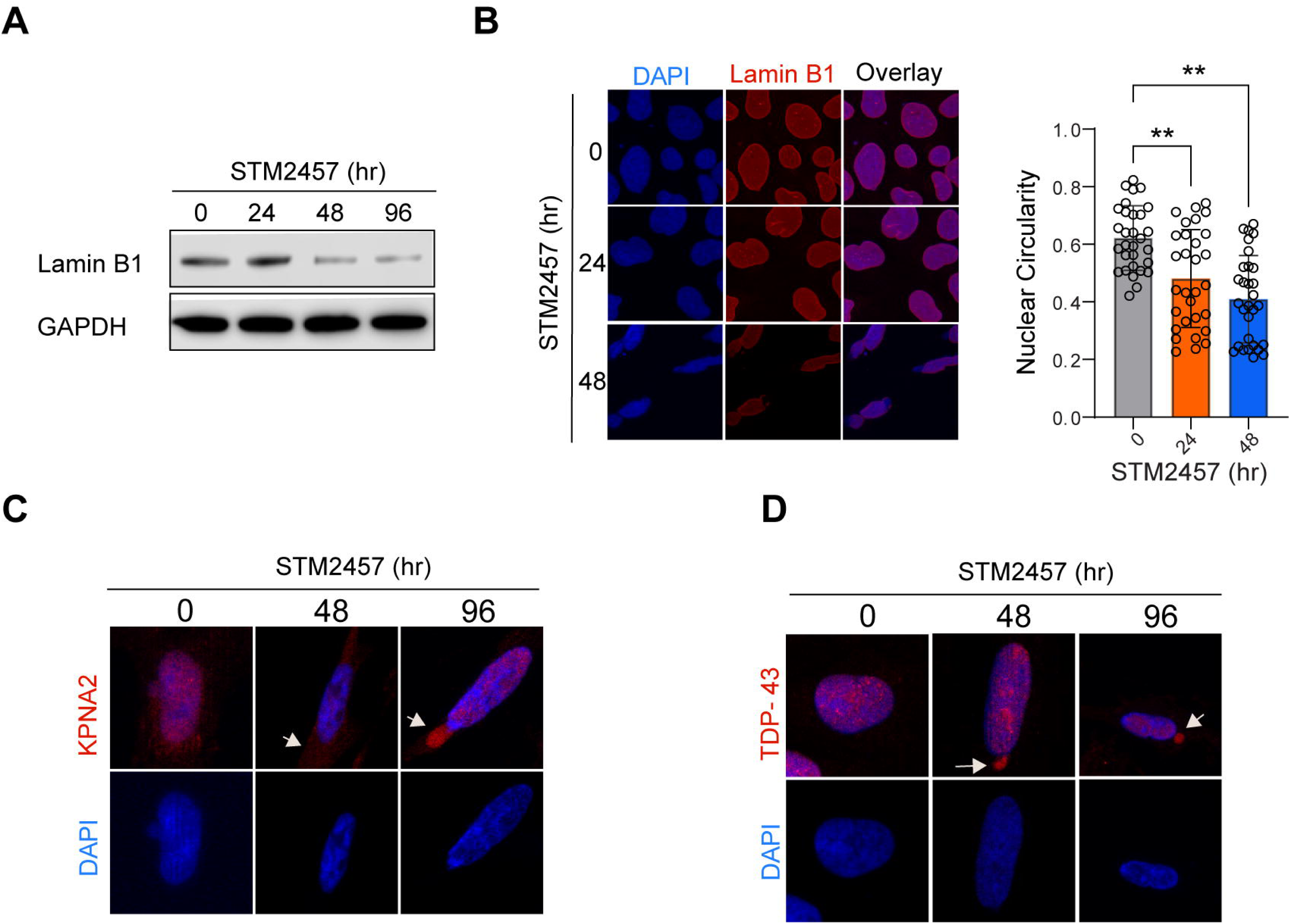
m6A inhibition remodels nuclear architecture and disrupts nuclear protein compartmentalization. (a) Representative western blot of Lamin B1 (LMNB1) protein levels in vehicle- and STM2457-treated HMC3 cells. (b) Representative LMNB1 immunofluorescence images and quantification of nuclear circularity in vehicle- and STM2457-treated cells. (c) Representative KPNA2 immunofluorescence showing redistribution of KPNA2 following STM2457 treatment. (d) Representative TDP-43 immunofluorescence showing nuclear localization in vehicle-treated cells and partial cytoplasmic mislocalization after STM2457 treatment. Data are presented as mean ± SEM; n = 3 independent biological experiments. Statistical significance was determined by two-tailed unpaired Student’s t test.

We next examined TDP-43, an RNA-binding protein that is normally enriched in the nucleus and whose cytoplasmic mislocalization is strongly associated with neurodegenerative disease. In vehicle-treated HMC3 cells, TDP-43 was predominantly nuclear. After 48 and 96 h of STM2457 treatment, a subset of cells showed increased cytoplasmic TDP-43 signal, indicating partial nuclear-to-cytoplasmic mislocalization (Figure 4d). These findings suggest that m6A inhibition is accompanied by altered nuclear compartmentalization of RNA-binding and nuclear transport-associated proteins.

Together, reduced nuclear envelope and nuclear pore gene expression, LMNB1 protein loss, altered nuclear morphology, KPNA2 redistribution, and TDP-43 mislocalization indicate that nuclear architectural remodeling is an underappreciated feature of the m6A inhibition-induced senescence-like state in microglia. These findings extend the phenotype beyond inflammatory activation and cell-cycle arrest, suggesting that reduced m6A also disrupts the structural organization and protein compartmentalization of the nucleus.

## Discussion

Our study identifies m6A as a safeguard against senescence-like inflammatory remodeling in human microglia. Selective inhibition of METTL3 catalysis was sufficient to induce multiple features of senescence, including increased SA-β-gal activity, altered cellular morphology, proliferative arrest, suppression of mitotic gene programs, CDKN2A/p16 induction, Lamin B1 loss, and disruption of nuclear architecture. These changes occurred together with a strong NF-κB-and interferon-associated inflammatory program, supporting a model in which m6A helps maintain microglial homeostasis and restrains transition into a persistent inflammatory state with key hallmarks of senescence.

A central strength of this work is that it separates catalytic m6A loss from depletion of the METTL3/METTL14 writer complex. Earlier studies linking METTL3 or METTL14 to senescence have largely relied on genetic depletion of writer-complex subunits, which removes both m6A deposition and methylation-independent functions of these proteins. By using STM2457 to inhibit METTL3 catalysis while leaving the writer complex intact, our study shows that reduction of the m6A mark itself can initiate inflammatory and structural remodeling. Thus, m6A loss is not simply a passive correlate of senescence, but an instructive event capable of driving a senescence-like microglial state.

These findings also help reconcile apparently divergent roles of METTL3 in microglia. Prior work in acute injury and stimulation models has positioned METTL3 as a positive regulator of microglial activation, with METTL3 inhibition or depletion dampening inflammatory responses^21^. Our results do not necessarily contradict these studies. Instead, they suggest that m6A has state-dependent functions. During acute activation, m6A may support rapid inflammatory transcript production, processing, or stability. Sustained reduction of m6A over days appears to produce a distinct state marked by proliferative arrest, nuclear remodeling, cytoplasmic dsRNA accumulation, and SASP-like inflammatory signaling. This distinction between acute activation and chronic senescence-associated inflammation may explain why m6A inhibition appears anti-inflammatory in some microglial injury models but pro-inflammatory in our system.

How m6A loss engages inflammatory signaling remains an important question. Since m6A has been implicated in the control of repeat-derived RNAs, and senescent cells can derepress repetitive elements and accumulate immunostimulatory RNA, we initially considered broad transposable-element activation as a potential trigger. Our 48-h transcriptomic data do not support this model. Repeat-aware analyses showed that transposable-element changes were limited and selective, with no global induction of LINE, SINE, or LTR families. Thus, the increase in cytoplasmic dsRNA and the accompanying inflammatory program should not be interpreted as evidence of genome-wide transposable-element derepression.

Instead, our findings support a more selective model of immunostimulatory RNA accumulation after m6A loss. STM2457 induced robust cytoplasmic dsRNA accumulation and selectively upregulated HERVK-related transcripts, including the internal repeat annotation HERVK3-int. This HERVK-associated signal may be biologically meaningful, as ERV reactivation during aging has been shown to reinforce cellular senescence and tissue aging, with HERVK (HML-2) contributing to senescence-associated phenotypes^17^. Together, these findings suggest that m6A loss can engage selected ERV-associated features of senescence without requiring broad transposable-element derepression. They also indicate that cytoplasmic dsRNA accumulation in this setting is not simply a consequence of genome-wide repeat activation.

Nuclear remodeling represents another major component of the m6A-loss phenotype. STM2457 reduced Lamin B1, altered nuclear morphology, and disrupted nuclear speckles. Lamin B1 loss is a well-established marker of senescence and reflects changes in nuclear-envelope and chromatin organization. Nuclear speckles are RNA-processing hubs in which components of the m6A machinery are enriched^34^. The co-occurrence of reduced m6A, altered nuclear speckles, Lamin B1 loss, and inflammatory gene expression suggests that m6A loss affects not only transcript abundance but also the nuclear RNA-processing environment. Whether these nuclear changes contribute directly to immunogenic RNA formation or arise as part of the broader senescence-like program remains to be determined.

These observations may have broader relevance for brain aging. Microglial aging is marked by chronic inflammatory signaling, impaired surveillance, altered metabolic states, and increased vulnerability to neurodegenerative stress. Our findings suggest that disruption of an epitranscriptomic pathway can push human microglia toward an inflammatory senescence-like state. This raises the possibility that age-associated changes in RNA modification contribute to maladaptive microglial states in the aging brain.

In summary, catalytic reduction of m6A is sufficient to induce senescence-like inflammatory and nuclear remodeling in human microglia. Rather than causing broad transposable-element derepression, m6A loss produces a selective RNA response characterized by cytoplasmic dsRNA accumulation and HERVK-associated changes, together with Lamin B1 loss, altered nuclear morphology, and disruption of nuclear protein compartmentalization. These findings identify m6A as a safeguard against immunogenic RNA accumulation and nuclear architectural remodeling, and suggest that reduced m6A-mediated RNA regulation may contribute to chronic microglial inflammatory states during brain aging.

## Experimental procedures

### Cell culture

HMC3 human microglial cells (ATCC, CRL-3304) were maintained at 37°C in a humidified incubator with 5% CO_2_ in high-glucose DMEM (Gibco, 11965118) supplemented with 10% fetal bovine serum (Gibco, 10437028) and 1% penicillin-streptomycin (Gibco, 15140-122). Cells were passaged before reaching confluence, used within 10 passages after thawing, and routinely monitored for morphology.

### STM2457 treatment

STM2457 was used to inhibit METTL3 catalytic activity and reduce global m6A. A 10 mM stock in DMSO was stored at -80 °C. HMC3 cells were treated with 20 µM STM2457 or an equivalent volume of DMSO vehicle, with a final DMSO concentration of 0.1% in all treatment and vehicle-control wells. Cells were collected or fixed after 24, 48 or 96 h of treatment, as indicated for each assay. Unless otherwise stated, experiments used three independent biological replicates per condition.

### RNA dot blot for global m6A

Total RNA was isolated with TRI Reagent (Zymo Research, R2050-1-200) and purified with the RNA Clean & Concentrator kit (Zymo Research, R1015). Equal amounts of RNA per sample were denatured at 85 °C for 5 min and spotted onto an Amersham Hybond-N+ membrane. After UV crosslinking, membranes were blocked in 3% BSA and incubated with anti-m6A antibody (Abcam, ab151230; 1:1000) overnight at 4 °C, followed by HRP-conjugated secondary antibody (Santa Cruz Biotechnology, sc-2357; 1:5000) and enhanced chemiluminescent substrate (LI-COR, 926-95000). Total RNA loading was assessed by methylene blue staining (Acros Organics, 41424-0250). Dot intensity was quantified in ImageJ/Fiji and normalized to total RNA signal.

### Cell staining and fluorescence imaging assays

HMC3 cells were seeded on sterile coverslips and treated with STM2457 or vehicle for the indicated times. Unless otherwise specified, cells were washed with PBS or DPBS, fixed with 4% paraformaldehyde in PBS for 15 min at room temperature, and mounted in Fluoromount-G (Thermo Fisher Scientific, 00-4958-02). All staining procedures were performed according to the manufacturers’ instructions where applicable.

For senescence-associated β-galactosidase (SA-β-gal) staining, cells were processed using the Cellular Senescence Detection Kit (SG04; Dojindo Molecular Technologies, Rockville, MD, USA). Vehicle- and STM2457-treated cells were washed with PBS, fixed with 4% paraformaldehyde, and incubated with SA-β-gal staining solution at 37 °C in the absence of CO₂ for 30 min.

Cell proliferation was assessed by EdU incorporation. Cells were incubated with 10 µM EdU for 2 h before fixation, and EdU-positive nuclei were detected using the Click-iT Plus EdU Cell Proliferation Kit (Thermo Fisher Scientific, C10639). Nuclei were counterstained with DAPI.

Lipid droplet was assessed using BODIPY 493/503. HMC3 cells were treated with STM2457 or vehicle for 0, 24, 48, or 96 h. At each time point, cells were incubated with 1 µM BODIPY 493/503 and 1 µM Hoechst 33342 in culture medium for 30 min at 37 °C protected from light. Cells were then washed twice with DPBS (Gibco, 14190-144), fixed with 4% paraformaldehyde, washed twice with PBS, and mounted.

For cell morphology analysis, F-actin was visualized by phalloidin staining. Cells grown on coverslips were washed twice with DPBS, fixed with 4% paraformaldehyde, permeabilized with 0.1% Triton X-100 in PBS for 15 min, and blocked with 3% BSA in PBS for 30 min at room temperature. Cells were stained with Phalloidin-iFluor 594 (Abcam, ab176757; 1:1000) and DAPI for 20 min at room temperature in the dark.

For general immunofluorescence staining, fixed cells were permeabilized with 0.1% Triton X-100 in PBS for 15 min and blocked with 3% normal donkey serum in PBS for 1 h at room temperature. Coverslips were incubated overnight at 4 °C with primary antibodies against dsRNA (J2 clone; EMD Millipore, MABE1134; 1:1000), HERV-K Env (AMSBIO, HERM-1811-5; 1:1000),

Lamin B1 (Proteintech, 12987-1-AP; 1:1000), or SRRM2 (Proteintech, 30741-1-AP; 1:1000). After washing, coverslips were incubated with Alexa Fluor-conjugated secondary antibodies for 1 h at room temperature, washed, and mounted in Fluoromount-G.

### Nuclear morphology and circularity analysis

Nuclear morphology was analyzed from Lamin B1 immunofluorescence images. Vehicle- and STM2457-treated HMC3 cells were fixed, immunostained for Lamin B1, and counterstained with DAPI as described above. Images were acquired under identical exposure and acquisition settings across conditions. Nuclear boundaries were segmented in Fiji/ImageJ using the DAPI channel, with Lamin B1 staining used to confirm nuclear-envelope labeling and exclude poorly segmented or overlapping nuclei. Nuclear circularity was calculated as described before^35^. Briefly, for each segmented nucleus in Fiji using the formula: circularity = 4π × area/perimeter². Circularity values range from 0 to 1, with a value of 1 indicating a perfectly circular nucleus and lower values indicating increasingly elongated or irregular nuclear shape.

### Western blotting

Cells were lysed in RIPA buffer (Thermo Fisher Scientific, 89901) supplemented with protease and phosphatase inhibitors (Roche Diagnostics). Protein concentration was determined by Bradford assay (Bio-Rad, 5000006). Equal amounts of protein (30 µg per sample) were separated by SDS-PAGE on NuPAGE 4–12% Bis-Tris gels (1.5 mm, 10-well; Invitrogen, NP0335BOX) and transferred to nitrocellulose membranes (Bio-Rad, 1620115). Membranes were blocked in TBST containing 5% skim milk and incubated overnight at 4 °C with primary antibodies against Lamin B1 (Proteintech, 12987-1-AP; 1:1000), METTL3 (Abcam, ab195352; 1:1000), METTL14 (Proteintech, 26158-1-AP; 1:1000) and GAPDH (Santa Cruz Biotechnology, sc-47724; 1:1000).

Membranes were washed and incubated for 1 h with HRP-conjugated secondary antibodies. Signals were detected with enhanced chemiluminescent substrate (LI-COR, 926-95000) on a LI-COR imaging system and quantified in ImageJ. Target protein levels were normalized to GAPDH.

### RNA extraction, RNA sequencing, and differential expression analysis

Total RNA was extracted from vehicle- and STM2457-treated HMC3 cells using TRI Reagent (Zymo Research, R2050-1-200) and purified with the RNA Clean & Concentrator kit (Zymo Research) according to the manufacturer’s instructions. Ribosomal RNA was depleted using the NEBNext rRNA Depletion Kit v2 for Human/Mouse/Rat samples (New England Biolabs, E7400/E7405). RNA-seq libraries were prepared from rRNA-depleted RNA and sequenced as paired-end reads on an Illumina NextSeq 1000/2000 system.

Paired-end reads were aligned to the human hg38 reference genome using STAR v2.7.10a. STAR was run to generate coordinate-sorted BAM files and gene-level read counts using the GeneCounts option. To retain reads informative for repetitive and multi-mapping regions, alignments were allowed to map to up to 20 loci using --outFilterMultimapNmax 20. Splice-junction filtering was performed using --outFilterType BySJout, --alignSJoverhangMin 8, - -alignSJDBoverhangMin 1, and --alignIntronMax 1000000.

Gene-level raw counts were analyzed in R using DESeq2. Differential expression was assessed between STM2457- and vehicle-treated cells. Genes with a Benjamini-Hochberg-adjusted P value < 0.05 and an absolute fold change ≥ 1.5 were considered differentially expressed for threshold-based summaries and visualization. Variance-stabilized counts were used for sample-level quality control, principal-component analysis, and heatmap visualization.

### Transposable-element and repeat-family analysis

Repeat-derived transcript abundance was quantified from RNA-seq alignments using the repeat-aware tool Telescope and TEtranscripts. For the primary repeat-family analysis, LINE, LTR, and SINE features were quantified separately using RepeatMasker-derived annotations, and the resulting count matrices were used for downstream differential expression analysis.

Differential repeat expression was analyzed independently for LINE, LTR, and SINE features using DESeq2. To avoid estimating normalization factors directly from repeat-derived counts, DESeq2 size factors were first estimated from the matched gene-level raw count matrix and then applied to each repeat-feature count matrix. Gene counts and repeat features with fewer than 10 total counts across all samples were excluded before analysis. Count values generated from repeat quantification were rounded to integers before DESeq2 analysis.

### Image analysis and quantification

Image analyses were performed using ImageJ/Fiji unless otherwise stated. For each experiment, images were acquired using identical exposure, laser power, detector gain, and acquisition settings across conditions. Segmentation parameters and intensity thresholds were kept constant within each experiment. Quantification was performed across independent biological replicates, with the number of replicates, fields, and cells reported in the corresponding figure legends.

For cell-area measurements, individual cell boundaries were segmented from phase-contrast or fluorescence images, and cell area was quantified for each segmented cell. For fluorescence-intensity measurements, background-subtracted mean fluorescence intensity was calculated for each cell or nucleus, as appropriate for the marker analyzed. Cell-level or field-level measurements were averaged within each biological replicate before statistical analysis.

### Statistical analysis

Statistical analyses were performed in R unless otherwise stated. For comparisons between two groups, an unpaired two-tailed Student’s t test was used when data met parametric assumptions. For comparisons involving multiple treatment groups or time points, one-way or two-way ANOVA followed by Tukey’s multiple-comparisons test was used as appropriate. RNA-seq and transposable-element differential-expression analyses were corrected for multiple testing using the Benjamini-Hochberg method. Fisher’s exact test was used to evaluate overlap between gene lists where indicated. Data are presented as mean ± SEM unless otherwise stated. The number of biological replicates, quantified cells or fields, and the statistical tests used are reported in the figure legends.

## Acknowledgements

We thank the Cancer Prevention and Research Institute of Texas (CPRIT, RP180734) for sequencing support, and members of the Weng laboratory for critical review of the manuscript.

## Confit of interest

The authors declare no conflict of interest.

## Author contributions

D.P. and Y.-L.W. conceived and designed the study. R.B. constructed the RNA-sequencing libraries. D.P. and R.B. performed the experiments and collected the data. L.S. performed the RNA-seq analysis. Y.-L.W. supervised the study and acquired funding. All authors contributed to editing and approved the final version of the manuscript.

## Funding information

This study was supported by the National Institutes of Health (R01ES031511 to Y.-L.W.).

## Data availability

All data supporting the findings are available from the corresponding author upon reasonable request.

## Notes

### Competing Interest Statement

The authors have declared no competing interest.

## References

1 Liu, K. et al. Senescent Microglia Mediate Neuroinflammation-Induced Cognitive Dysfunction by Selective Elimination of Excitatory Synapses in the Hippocampal CA1. Aging Cell 24, e70167, doi:10.1111/acel.70167 (2025).

2 Rim, C., You, M. J., Nahm, M. & Kwon, M. S. Emerging role of senescent microglia in brain aging-related neurodegenerative diseases. Transl Neurodegener 13, 10, doi:10.1186/s40035-024-00402-3 (2024).

3 Matsudaira, T. et al. Cellular senescence in white matter microglia is induced during ageing in mice and exacerbates the neuroinflammatory phenotype. Commun Biol 6, 665, doi:10.1038/s42003-023-05027-2 (2023).

4 Rachmian, N. et al. Identification of senescent, TREM2-expressing microglia in aging and Alzheimer’s disease model mouse brain. Nat Neurosci 27, 1116–1124, doi:10.1038/s41593-024-01620-8 (2024).

5 Ocanas, S. R. et al. Microglial senescence contributes to female-biased neuroinflammation in the aging mouse hippocampus: implications for Alzheimer’s disease. J Neuroinflammation 20, 188, doi:10.1186/s12974-023-02870-2 (2023).

6 Meyer, K. D. et al. Comprehensive analysis of mRNA methylation reveals enrichment in 3’ UTRs and near stop codons. Cell 149, 1635–1646, doi:10.1016/j.cell.2012.05.003 (2012).

7 Liu, J. et al. A METTL3-METTL14 complex mediates mammalian nuclear RNA N6-adenosine methylation. Nat Chem Biol 10, 93–95, doi:10.1038/nchembio.1432 (2014).

8 Yue, Y., Liu, J. & He, C. RNA N6-methyladenosine methylation in post-transcriptional gene expression regulation. Genes Dev 29, 1343–1355, doi:10.1101/gad.262766.115 (2015).

9 Chelmicki, T. et al. m(6)A RNA methylation regulates the fate of endogenous retroviruses. Nature 591, 312–316, doi:10.1038/s41586-020-03135-1 (2021).

10 Zhang, W. et al. METTL3-dependent m6A RNA methylation regulates transposable elements and represses human naive pluripotency through transposable element-derived enhancers. Nucleic Acids Res 53, doi:10.1093/nar/gkaf349 (2025).

11 Barter, B. & Cho, J. RNA methylation in retrotransposon control. Trends Genet 41, 556–558, doi:10.1016/j.tig.2025.04.013 (2025).

12 Roulois, D. et al. DNA-Demethylating Agents Target Colorectal Cancer Cells by Inducing Viral Mimicry by Endogenous Transcripts. Cell 162, 961–973, doi:10.1016/j.cell.2015.07.056 (2015).

13 Ahmad, S. et al. Breaching Self-Tolerance to Alu Duplex RNA Underlies MDA5-Mediated Inflammation. Cell 172, 797–810 e713, doi:10.1016/j.cell.2017.12.016 (2018).

14 Chiappinelli, K. B. et al. Inhibiting DNA Methylation Causes an Interferon Response in Cancer via dsRNA Including Endogenous Retroviruses. Cell 162, 974–986, doi:10.1016/j.cell.2015.07.011 (2015).

15 Rehwinkel, J. & Gack, M. U. RIG-I-like receptors: their regulation and roles in RNA sensing. Nat Rev Immunol 20, 537–551, doi:10.1038/s41577-020-0288-3 (2020).

16 De Cecco, M. et al. L1 drives IFN in senescent cells and promotes age-associated inflammation. Nature 566, 73–78, doi:10.1038/s41586-018-0784-9 (2019).

17 Liu, X. et al. Resurrection of endogenous retroviruses during aging reinforces senescence. Cell 186, 287–304 e226, doi:10.1016/j.cell.2022.12.017 (2023).

18 Wu, Z. et al. METTL3 counteracts premature aging via m6A-dependent stabilization of MIS12 mRNA. Nucleic Acids Res 48, 11083–11096, doi:10.1093/nar/gkaa816 (2020).

19 Li, Q. et al. NSUN2-Mediated m5C Methylation and METTL3/METTL14-Mediated m6A Methylation Cooperatively Enhance p21 Translation. J Cell Biochem 118, 2587–2598, doi:10.1002/jcb.25957 (2017).

20 Liu, P. et al. m(6)A-independent genome-wide METTL3 and METTL14 redistribution drives the senescence-associated secretory phenotype. Nat Cell Biol 23, 355–365, doi:10.1038/s41556-021-00656-3 (2021).

21 Wen, L. et al. The m6A methyltransferase METTL3 promotes LPS-induced microglia inflammation through TRAF6/NF-kappaB pathway. Neuroreport 33, 243–251, doi:10.1097/WNR.0000000000001550 (2022).

22 Wu, X. et al. The m(6)A methyltransferase METTL3 drives neuroinflammation and neurotoxicity through stabilizing BATF mRNA in microglia. Cell Death Differ 32, 100–117, doi:10.1038/s41418-024-01329-y (2025).

23 Yankova, E. et al. Small-molecule inhibition of METTL3 as a strategy against myeloid leukaemia. Nature 593, 597–601, doi:10.1038/s41586-021-03536-w (2021).

24 Marschallinger, J. et al. Lipid-droplet-accumulating microglia represent a dysfunctional and proinflammatory state in the aging brain. Nat Neurosci 23, 194–208, doi:10.1038/s41593-019-0566-1 (2020).

25 Claes, C. et al. Plaque-associated human microglia accumulate lipid droplets in a chimeric model of Alzheimer’s disease. Mol Neurodegener 16, 50, doi:10.1186/s13024-021-00473-0 (2021).

26 Coppe, J. P. et al. Senescence-associated secretory phenotypes reveal cell-nonautonomous functions of oncogenic RAS and the p53 tumor suppressor. PLoS Biol 6, 2853–2868, doi:10.1371/journal.pbio.0060301 (2008).

27 Coppe, J. P., Desprez, P. Y., Krtolica, A. & Campisi, J. The senescence-associated secretory phenotype: the dark side of tumor suppression. Annu Rev Pathol 5, 99–118, doi:10.1146/annurev-pathol-121808-102144 (2010).

28 Yoneyama, M. et al. The RNA helicase RIG-I has an essential function in double-stranded RNA-induced innate antiviral responses. Nat Immunol 5, 730–737, doi:10.1038/ni1087 (2004).

29 Kato, H. et al. Differential roles of MDA5 and RIG-I helicases in the recognition of RNA viruses. Nature 441, 101–105, doi:10.1038/nature04734 (2006).

30 Bendall, M. L. et al. Telescope: Characterization of the retrotranscriptome by accurate estimation of transposable element expression. PLoS Comput Biol 15, e1006453, doi:10.1371/journal.pcbi.1006453 (2019).

31 Jin, Y., Tam, O. H., Paniagua, E. & Hammell, M. TEtranscripts: a package for including transposable elements in differential expression analysis of RNA-seq datasets. Bioinformatics 31, 3593–3599, doi:10.1093/bioinformatics/btv422 (2015).

32 Freund, A., Laberge, R. M., Demaria, M. & Campisi, J. Lamin B1 loss is a senescence-associated biomarker. Mol Biol Cell 23, 2066–2075, doi:10.1091/mbc.E11-10-0884 (2012).

33 Shah, P. P. et al. Lamin B1 depletion in senescent cells triggers large-scale changes in gene expression and the chromatin landscape. Genes Dev 27, 1787–1799, doi:10.1101/gad.223834.113 (2013).

34 Ping, X. L. et al. Mammalian WTAP is a regulatory subunit of the RNA N6-methyladenosine methyltransferase. Cell Res 24, 177–189, doi:10.1038/cr.2014.3 (2014).

35 Matias, I. et al. Loss of lamin-B1 and defective nuclear morphology are hallmarks of astrocyte senescence in vitro and in the aging human hippocampus. Aging Cell 21, e13521, doi:10.1111/acel.13521 (2022).

